# Synaptic vesicle glycoprotein 2C enhances vesicular storage of dopamine and counters dopaminergic toxicity

**DOI:** 10.1101/2023.06.26.546143

**Authors:** Meghan L Bucher, Amy R Dunn, Joshua M Bradner, Kristen Stout Egerton, James P Burkett, Michelle A Johnson, Gary W Miller

**Affiliations:** Department of Environmental Health Sciences, Mailman School of Public Health, Columbia University, New York, NY 10032, USA; Department of Environmental Health, Rollins School of Public Health, Emory University, Atlanta, GA 30322, USA; Department of Molecular Pharmacology and Therapeutics, Vagelos College of Physicians and Surgeons, Columbia University, New York, NY 10031, USA

## Abstract

Dopaminergic neurons of the substantia nigra exist in a persistent state of vulnerability resulting from high baseline oxidative stress, high energy demand, and broad unmyelinated axonal arborizations. Impairments in the storage of dopamine compound this stress due to cytosolic reactions that transform the vital neurotransmitter into an endogenous neurotoxicant, and this toxicity is thought to contribute to the dopamine neuron degeneration that occurs Parkinson’s disease. We have previously identified synaptic vesicle glycoprotein 2C (SV2C) as a modifier of vesicular dopamine function, demonstrating that genetic ablation of SV2C in mice results in decreased dopamine content and evoked dopamine release in the striatum. Here, we adapted a previously published *in vitro* assay utilizing false fluorescent neurotransmitter 206 (FFN206) to visualize how SV2C regulates vesicular dopamine dynamics and determined that SV2C promotes the uptake and retention of FFN206 within vesicles. In addition, we present data indicating that SV2C enhances the retention of dopamine in the vesicular compartment with radiolabeled dopamine in vesicles isolated from immortalized cells and from mouse brain. Further, we demonstrate that SV2C enhances the ability of vesicles to store the neurotoxicant 1-methyl-4-phenylpyridinium (MPP^+^) and that genetic ablation of SV2C results in enhanced 1-methyl-4-phenyl-1,2,3,6-tetrahydropyridine (MPTP)-induced vulnerability in mice. Together, these findings suggest that SV2C functions to enhance vesicular storage of dopamine and neurotoxicants, and helps maintain the integrity of dopaminergic neurons.

## Introduction

Deficiency in dopamine neurotransmission, resulting from dysfunction and degeneration of dopaminergic neurons in the substantia nigra, is the key pathologic feature of Parkinson’s disease.^1^ Effective dopaminergic neurotransmission regulates motor function and depends on proper dopamine homeostasis including cytosolic synthesis, vesicular packaging, evoked synaptic release, post-synaptic receptor activation, pre-synaptic reuptake, and enzymatic metabolism.^2^ Dysregulation of dopamine homeostasis can cause deficits in dopaminergic neurotransmission and jeopardize neuronal health. At baseline, cytosolic dopamine that undergoes enzymatic catabolism by monoamine oxidase generates electrons that are utilized by the electron transport chain in mitochondria to promote energy production and regulate neuronal activity.^3^ However, increased neuronal activity or external stressors can overwhelm the endogenous antioxidant defense system leading dopamine to act as an endogenous neurotoxin that generates highly reactive metabolites such as dopamine quinone, toxic aldehydes such as DOPAL, and reactive oxygen species.^4,5^ These reactive products can contribute to cellular dysfunction through interactions with proteins, lipids, and nucleic acids.^3,6^ The generation of these toxic species further compounds the inherent sensitivity of dopamine neurons which typically have high energy demands and broad, unmyelinated axons with large terminal fields that have high metabolic requirements.^7-10^

Maintaining a low amount of cytosolic dopamine is crucial to cellular health. The vesicular monoamine transporter 2 (VMAT2) minimizes the cytosolic pool of dopamine by sequestering dopamine into synaptic vesicles. There is significant evidence that dysregulation of dopamine homeostasis can cause dopaminergic neurons to become vulnerable to dysfunction and degeneration, and replicate features of Parkinson’s disease.^11-14^ There are several documented cases of infantile parkinsonism resorting from variants in *SLC18A2* – the gene for VMAT2 – that are believed to result from decreased VMAT2 expression or function.^15,16^ Conversely, rare SNPs in *SLC18A2*, which are believed to be gain-of-function mutations resulting in enhanced VMAT2 expression or function, confer a decrease in the risk of developing Parkinson’s disease, and two low-activity variants in VMAT2 have been identified that may be associated with an increased risk of developing the disease.^17-20^ Furthermore, post-mortem tissue from Parkinson’s disease patients demonstrates decreased VMAT2 immunoreactivity, and analysis of vesicles isolated from post-mortem brain tissue demonstrate decreased VMAT2-mediated dopamine uptake in Parkinson’s disease patients.^21,22^ The impairments in VMAT2 are not limited to the brain, as analysis of circulating platelets in Parkinson’s disease patients demonstrated decreased VMAT2 mRNA suggesting a systemic deficiency in VMAT2.^23^ However, even with functioning VMAT2, vesicles are inherently permeant and allow dopamine to passively leak into the cytoplasm; thus, it is important to evaluate other factors that contribute to the cytosolic pool of dopamine such as factors that regulate the retention of dopamine within vesicles.^24^

Recent investigation has focused on additional factors that regulate vesicular dynamics of dopamine, both at the basic biological level and for disease relevance. Although it is unclear which vesicular components may contribute to vesicular leakage, several lines of evidence suggest that synaptic vesicle glycoproteins (SV2s) may be capable of facilitating the storage of intravesicular molecules, including neurotransmitters, due to a heavily glycosylated intraluminal loop.^25-27^ SV2 proteins, of which there are three paralogues (SV2A, SV2B, and SV2C), are synaptic vesicle localized glycoproteins in the SLC22B family of solute carriers.^28^ The three SV2s have differential expression throughout the brain, with SV2A and SV2B having more ubiquitous expression, and SV2C having enriched expression in the basal ganglia, particularly in dopaminergic neurons.^29^ While SV2 proteins are identified as solute carriers, the substrates remain unidentified. There is some evidence to suggest that SV2A is a galactose transporter, which corroborates analysis identifying sequence homology of SV2 proteins to bacterial proteins that transport sugars, but this has not been shown directly.^30,31^ SV2 proteins also appear to enable calcium-mediated exocytosis of synaptic vesicles, and it has further been hypothesized that SV2C may be a novel neurotransmitter transporter.^32-34^

SV2C was first implicated in Parkinson’s disease following identification in a genome-wide association study (GWAS) as a modifier of nicotine’s protective effect against developing Parkinson’s disease.^35^ Intriguingly, we identified that genetic ablation of SV2C alters the effect of nicotine on dopamine transmission, and a 2022 study investigating this relationship demonstrated the necessity of drosophila SV2 orthologs in nicotine-mediated protection against alpha-synuclein induced neurodegeneration.^36,37^ Furthermore, subsequent GWAS identified SV2C as a modifier of GBA-associated PD risk, Parkinson’s disease patient’s response to L-DOPA, and most recently directly as a risk-modifier for Parkinson’s disease.^38-41^ Additionally, our laboratory has demonstrated aberrant SV2C staining patterns in post-mortem brain tissue from people with Parkinson’s disease, and an interaction between SV2C and alpha-synuclein.^36^

Although we have not identified a direct protein-protein interaction between VMAT2 and SV2C, these proteins appear to work in concert to modulate dopamine dynamics.^42^ A study that performed transcriptomics in mice with knock-out of VMAT2 in norepinephrine neurons identified SV2C as the gene with the highest enrichment in expression following VMAT2 knock-out in multiple brain areas.^43^ We have additionally shown that animals lacking SV2C have decreased dopamine content in the striatum, as well as impaired evoked dopamine release.^36^ These animals also displayed decreased locomotor activity correlating with the decrease in dopamine transmission.^36^ To better understand the biological function of SV2C and its role mediating Parkinson’s disease risk, this work uses a range of pharmacological and toxicological approaches in cellular models, isolated vesicles, and *in vivo* studies in mice to demonstrate that SV2C controls vesicular retention of dopamine and ameliorates dopaminergic toxicity

## Methods

### Cell lines

Human embryonic kidney (HEK293) cells were stably transfected with human vesicular monoamine transporter 2 (VMAT2; HEK-VMAT2) utilizing zeocin selection.^44^ A secondary stable transfection was performed on the HEK-VMAT2 cell line to add human synaptic vesicle glycoprotein 2C (SV2C) expression (HEK-VMAT2-SV2C) utilizing geneticin selection. HEK-VMAT2 cells were maintained in media consisting of Dulbecco’s Modified Eagle Medium (DMEM) with 4.5g/L glucose (Corning), 10% fetal bovine serum (Fisher), 0.5% penicillin-streptomycin (Sigma), and 100mg/ml Zeocin (Fisher). HEK-VMAT2-SV2C cells were maintained in media consisting of DMEM with 4.5g/L glucose, 10% fetal bovine serum, 0.5% Penn/Strep, 100mg/ml Zeocin, and 250ug/ml geneticin (Fisher). All cells were maintained in an incubator at 37ºC with 5% CO_2_ on 10cm cell culture dishes coated with poly-D-Lysine (Sigma).

### Western blot

Western blots were performed as previously described.^36^ Cell culture samples were collected in RIPA buffer, sonicated, centrifuged and the nuclear fraction discarded, the cytoplasmic fraction was used for subsequent western analysis. Samples were subjected to SDS-PAGE and transferred to a nitrocellulose membrane. Membranes were blocked with 5% nonfat dry milk and incubated in primary antibody (SV2C, 1:1,000 (Fisher, 501733345); VMAT2, 1:2000 (Invitrogen, MA5-24939); ACTIN, 1:5,000 (Sigma, A5060); GAPDH, 1:5,000 (Invitrogen, PA1-9046)) overnight at 4°C with gentle agitation. Secondary antibodies (Fluorescent channels 700 and 800, 1:15,000 (Licor, IRDye 800cw and IRDye 680id (Fisher)) were incubated at room temperature for 1 hour. Signal was visualized using the Licor Odyssey system. Unilateral striatal dissections were homogenized and underwent differential centrifugation to achieve a crude synaptosomal protein preparation. 20μg of protein was run through an SDS-PAGE gel and transferred to a PVDF membrane. Nonspecific antibody binding was blocked with a 7.5% nonfat dry milk solution, and the membrane was incubated in primary antibody overnight at 4°C with gentle agitation. Membranes were then incubated in HRP-conjugated secondary antibody for 1 hour at room temperature. Protein was visualized using chemiluminescence (Thermo Fisher) and a BioRad UV imager. Protein was quantified using ImageLab software and normalized to an actin loading control. Rabbit anti-TH (1:1,000 (Millipore AB152)). For experiments using SV2C-KO mouse tissue, lack of SV2C protein expression was confirmed via immunoblotting, and samples with a genotype-protein expression mismatch were excluded from analysis.

### Immunocytochemistry

Cells were seeded onto a poly-D-Lysine (Sigma) coated chamber slide and allowed to attach overnight. To perform immunocytochemistry, media was removed and cells were fixed with 2% paraformaldehyde warmed to 37ºC for 5m. After 5m, cells were fixed with 4% paraformaldehyde at room temperature for 20m with gentle agitation. Cells were then rinsed with phosphate buffered saline (PBS (Fisher)) three times for 5m each before treatment with blocking solution (PBS with 10% normal goat serum (Fisher), 0.1% triton-x (Sigma), and 1% bovine serum albumin (Sigma)) at room temperature for 1h with gentle agitation. Following blocking, cells were treated with primary antibody solution (PBS with 10% normal goat serum, 0.1% triton-x, and 1% bovine serum albumin) with mouse anti-SV2C (1:500 (Sigma, MABN367)) and rabbit anti-VMAT2 (1:500 (Miller lab)) overnight at 4ºC with gentle agitation. After overnight incubation with primary antibodies, cells were washed in PBS with 0.1% triton three times for 5m each before treatment with secondary antibody solution (PBS with 1% normal goat serum, 0.1% Triton-x, and 1% bovine serum albumin) containing 1:400 goat anti-mouse Alexafluor 594 (Thermo Fisher) and 1:400 Goat anti-rabbit AlexaFluor 488 (Thermo Fisher) for 1h at room temperature with gentle agitation. Following secondary incubation, cells were washed with PBS three times for 5m each before coverslipping with Vectashield (Fisher). Microscopy images were acquired on an EVOS_*fl*_ microscope, and pseudo-colored in FIJI (ImageJ).

### FFN206 uptake assay

FFN206 uptake assay was performed as previously described.^44^ Briefly, HEK-VMAT2 and HEK-VMAT2-SV2C cells were seeded in DMEM + 0.5% P/S + 10% FBS in a black walled with clear flat bottom half-area 96-well plate (Corning) at 40,000 cells. 24h later, at confluency, culture media was removed and cells were pre-incubated in standard incubator conditions with DMSO or 1μM tetrabenazine for 30m in DMEM with 4.5g/L glucose, sodium pyruvate; without L-glutamine, phenol red (Corning). After 30m, FFN206 prepared in DMEM with 4.5g/L glucose, sodium pyruvate; without L-glutamine, phenol red (Corning) was added to a final concentration of 2μM and incubated for 1hr. After the incubation period, media was removed and replaced with DMEM with 4.5g/L glucose, sodium pyruvate; without L-glutamine, phenol red (Corning) containing 1:25 trypan blue and plate was scanned on BioTek Synergy H1 multiplate reader at 37ºC using endpoint fluorescent intensity measurements with excitation λ = 369 and excitation λ = 464. Fluorescent values were corrected by subtracting background fluorescent values obtained from cells on the same plate with same treatments without FFN206. For uptake experiments, each 96-well plate contained 12 technical replicates per cell line:treatment group. An average for the control condition (HEK-VMAT2:DMSO) was calculated and each well value was expressed as a percent of this control. A single averaged value was calculated for each cell line:treatment group to represent an experimental replicate. Two-way ANOVA with Šídák’s multiple comparisons test was performed in GraphPad Prism 9 on 7 experimental replicates. Tetrabenazine IC_50_ calculation was performed in GraphPad Prism 9 by non-linear regression ([inhibitor] vs. normalized response) and regressions were compared to determine significant differences. Data points were excluded only in cases of pipetting errors (e.g., well received no FFN solution). Microscopy images were acquired on an EVOS_*fl*_ microscope.

### FFN206 retention assay

HEK-VMAT2 and HEK-VMAT2-SV2C cells were seeded in DMEM + P/S + FBS in a black walled with clear flat bottom half-area 96-well plate (Corning) at 40,000 cells. 24h later, at confluency, culture media was removed and cells were incubated in standard culture incubator conditions with 2μM FFN206 in DMEM with 4.5g/L glucose, sodium pyruvate; without L-glutamine, phenol red (Corning) for 1hr. Following this 1hr incubation, media was removed and replaced with DMEM with 4.5g/L glucose, sodium pyruvate; without L-glutamine, phenol red (Corning) containing 1:25 trypan blue and plate was scanned on BioTek Synergy H1 multiplate reader at 37ºC using endpoint fluorescent intensity measurements with excitation λ = 369 and excitation λ = 464 for a baseline fluorescence value. DMSO (Fisher) or tetrabenazine (Sigma) were added for a final concentration of 0.02% DMSO and 1μM tetrabenazine. Fluorescent values were corrected by subtracting background fluorescent values obtained from cells on the same plate with same treatments without FFN206. For retention experiments, each 96-well plate contained 12 technical replicates per cell line:treatment group. Each well underwent background subtraction at each timepoint, and the fluorescent value was expressed as a percent of its own baseline fluorescent value at each timepoint. The percent of fluorescence was graphed over time and linear and non-linear regression (plateau followed by one-phase decay) analysis was performed in GraphPad Prism 9 to determine the rate of fluorescent decay and regressions were compared to determine significant differences. Data points were excluded only in cases of pipetting errors (e.g., well received no FFN solution). Data in repository features extended retention data to 60m.

### Vesicle isolation

Cells were grown to confluency in large (500cm^2^) cell culture dishes. Cells were collected in 20ml room temperature PBS, and pelleted (2 minutes at 1500 x g). After removing the supernatant, cells were re-suspended in 1200μl of ice-cold incomplete assay buffer (ICB) (100mM potassium tartrate, 25mM HEPES, 0.1mM EDTA, 0.05 mM EGTA, pH to 7.4), placed in a 3ml Dounce homogenizer and subjected to 30 strokes. Lysates were then run through the width of a 27-gauge needle twice for further homogenization. Lysates were centrifuged (8000 x g, 8 minutes, 4°C) and the supernatant collected as a crude vesicle preparation. Vesicles were further diluted in ice-cold ICB according to the number of samples needed for each experiment. A small aliquot was reserved for later BCA (Pierce) to determine protein concentration. All chemical reagents were from Sigma Aldrich. Brain-derived vesicles were collected from wild-type and SV2C-KO mice as previously described.^45^ Briefly, the cytoplasmic vesicular fraction was prepared from homogenized bilateral striata via differential centrifugation.

### Vesicular leakage

Crude vesicle fractions were isolated from HEK-VMAT2 cells and HEK-VMAT2-SV2C cells as described above. Ascorbate (1.7mM) and Mg^2+^ ATP salt (2mM) were added to ICB to make complete assay buffer (CAB). Dopamine hydrochloride (2μM) and 2% [^3^H]-dopamine (40nM) tracer were added to the CAB. 450μl of the CAB/dopamine solution was added to each glass test tube in the experiment and kept in racks at 30ºC in a water bath. Vesicles were isolated from WT and SV2C-KO mice, as described above. Samples were kept on ice and sequentially added to a 30ºC water bath. Cell culture vesicle samples were kept on ice and added sequentially in 50μl volumes to the CAB/dopamine tubes to begin the uptake reaction. Uptake proceeded for 10 minutes, at which time the VMAT2 inhibitor reserpine was added to the reactions to yield a final concentration of 10μM. Reserpine effectively blocks the transporter, preventing any further uptake of radiolabeled dopamine. Vesicular leak proceeded for 0, 5, 10, 15, and 20 minutes, with all time points staggered so that all samples concluded at the same time. After the “0” leak time point, cells were immediately harvested through a Brandel Cell Harvester onto GF/F filter paper coated with 0.5% polyethylenimine in ICB, and washed 3x with ice-cold ICB. Filter papers were cut, placed in scintillation vials and vortexed. Samples were stored in the dark and read 12 hours later using a liquid scintillation counter (Packard). A small portion of the crude vesicle lysate was analyzed by BCA (Pierce) analysis to normalize uptake to the amount of vesicles used and expressed as fmol of [^3^H] dopamine remaining per μg of vesicles. Each time point was replicated 4 times across 3 different experiments. VMAT2 specific uptake was calculated by subtracting the average uptake of vesicles pre-treated with reserpine. Experiments using radiolabeled MPP^+^ were performed in exactly the same way, substituting MPP^+^ (2μM) for dopamine hydrochloride and [^3^H] MPP^+^ for [^3^H] dopamine. Brain-derived vesicle samples were incubated for five minutes and then [^3^H] dopamine spiked in to a final concentration of 30nM. Uptake proceeded for 10 minutes, at which time VMAT2 inhibitor tetrabenazine was added to the samples to yield a final concentration of 10μM. This addition effectively froze the transporter, preventing additional uptake of radiolabeled dopamine. Vesicular leak was measured at 0, 1, 2.5, 5, 7.5, and 10 minutes. The experiment was arranged such that all samples concluded at the same time. Each time point was assessed in three specific and one nonspecific samples per animal. Nonspecific uptake, measured from samples that lacked ATP, was subtracted from total counts to generate specific uptake counts. Counts are shown as percentages of the zero leak time point.

Figures 4A and 6A represent total uptake without leak, or the “0” time point as an expression of maximal uptake. Unpaired t-tests were performed in GraphPad Prism 9 to determine differences in uptake. All chemical reagents were from Sigma Aldrich with the exception of radiolabeled compounds provided by Perkin Elmer. Non-linear regression (one-phase decay) was performed in GraphPad Prism 9 to determine the rate of radiolabeled decay and regressions were compared to determine significant differences. Data points were removed due to technical complications during experimental process or due to outlier exclusions based on values greater or less than one standard deviation from the mean.

**Figure 1.**
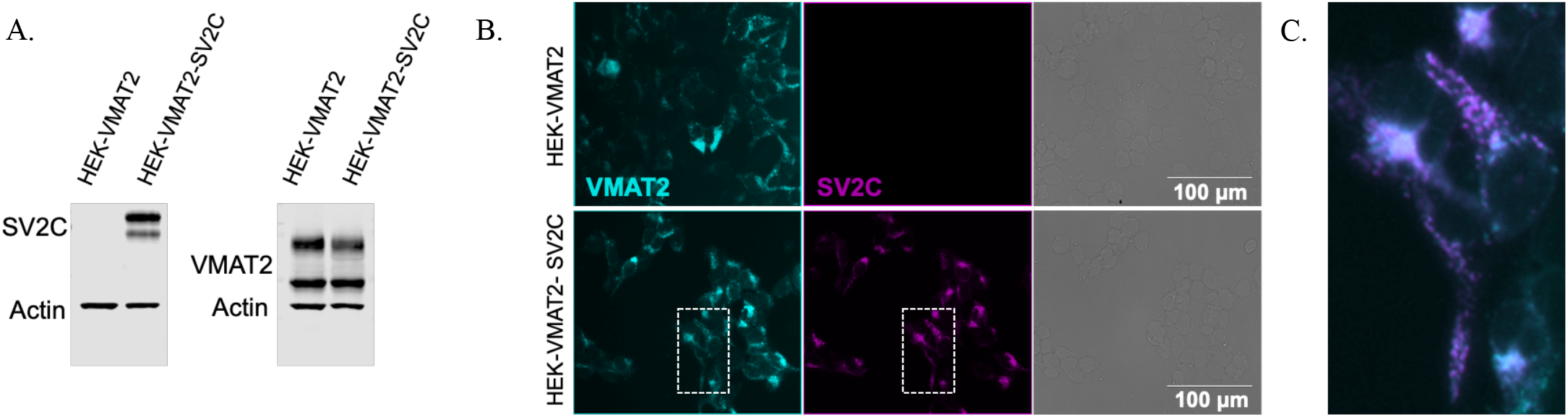
SV2C expression in HEK cells co-expressing VMAT2. Western blot and immunocytochemistry demonstrating SV2C expression in HEK-VMAT2-SV2C cells. A. Western blot performed on whole cell lysates to identify SV2C protein expression HEK-VMAT2-SV2C cells compared to HEK-VMAT2 cells lacking SV2C. B. Representative 40x immunocytochemistry images visualizing VMAT2 (cyan) and SV2C (magenta) protein expression in HEK-VMAT2 cells and HEK-VMAT2-SV2C cells (transmitted light with scale bar). C. SV2C co-localizes with VMAT2 in HEK-VMAT2-SV2C cells demonstrated by overlap of enlarged inset.

**Figure 2.**
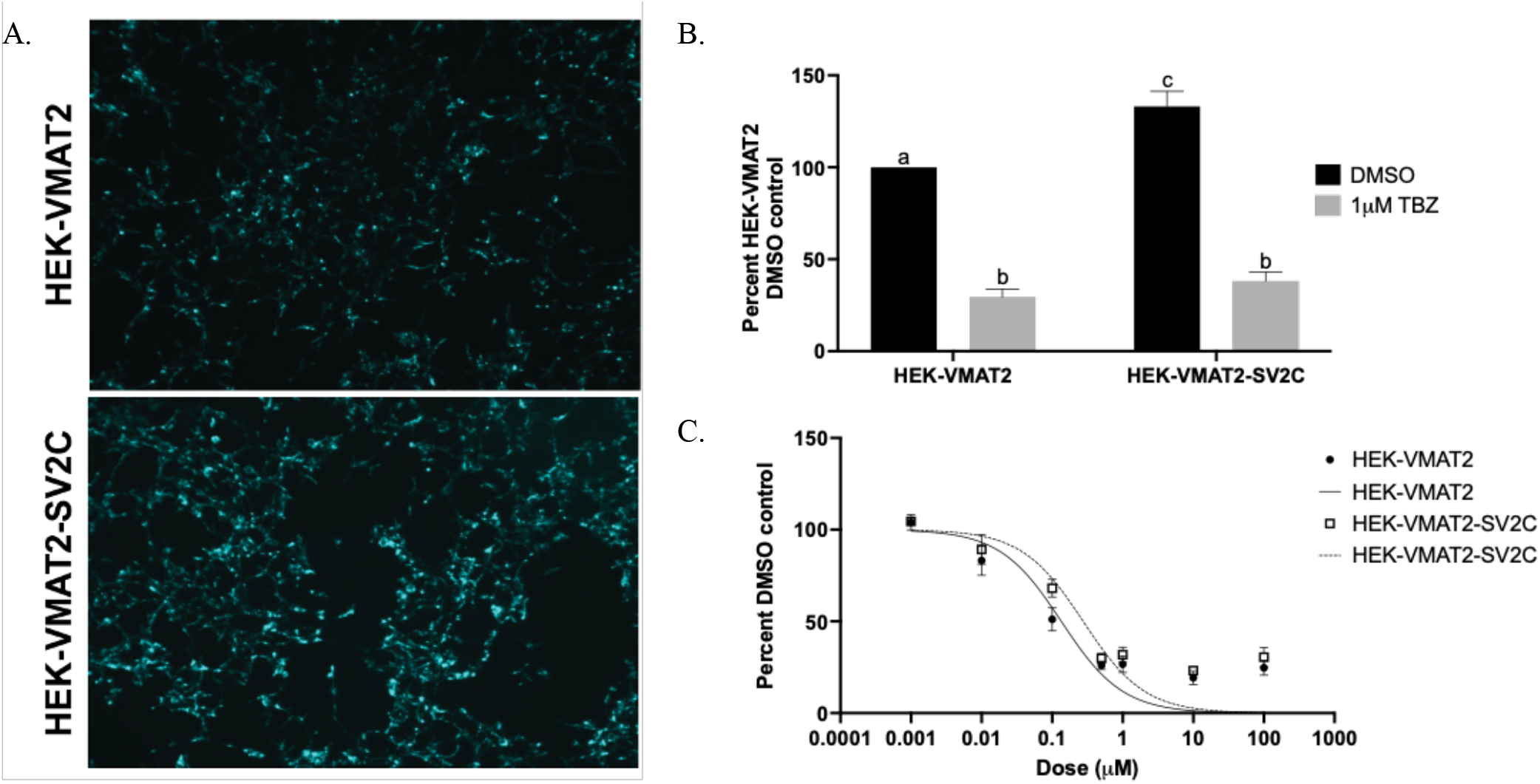
SV2C enhances vesicular uptake of the fluorescent dopamine analogue FFN206. A. Representative 10x images of FFN206 uptake in HEK-VMAT2 (top) and HEK-VMAT2-SV2C (bottom) cells. B. FFN206 uptake was measured in HEK-VMAT2 and HEK-VMAT2-SV2C cells seeded on 96-well plates following control (DMSO) or tetrabenazine (1μM) pretreatment. Values were transformed into percent control of the average HEK-VMAT2 DMSO value following background subtraction. For each experiment there were 12 technical replicates for each cell and treatment condition which were averaged to a single value for comparison across seven experimental replicates. Two-way ANOVA with Šídák’s multiple comparisons test. Effect of cell line (F(1, 24) = 15.49, p =0.0006); effect of treatment (F(1, 24) = 242.6, p <0.0001); effect of interaction (F(1, 24) = 5.207, p =0.0317), n = 7 experimental replicates, a:b p <0.0001, b:c p <0.0001, a:c p =0.0012. C. IC_50_ calculation for tetrabenazine in HEK-VMAT2 vs HEK-VMAT2-SV2C cells. FFN206 uptake was measured in HEK-VMAT2 and HEK-VMAT2-SV2C cells seeded on 96-well plates following control (DMSO) or tetrabenazine (0.001μM - 100μM) pretreatment. Values were transformed into percent control of the average HEK-VMAT2 DMSO value. For each experiment there were 4-6 technical replicates for each cell and treatment condition which were averaged to a single value for comparison across 4-8 experimental replicates. Non-linear regression to calculate IC_50_ values for each cell line demonstrated HEK-VMAT2 IC_50_ = 0.1348μM and HEK-VMAT2-SV2C IC_50_ = 0.2749μM. Comparison of non-linear regressions determined the curves of best fit were significantly different for each cell line (F(1, 102) = 5.469, p = 0.0213).

**Figure 3.**
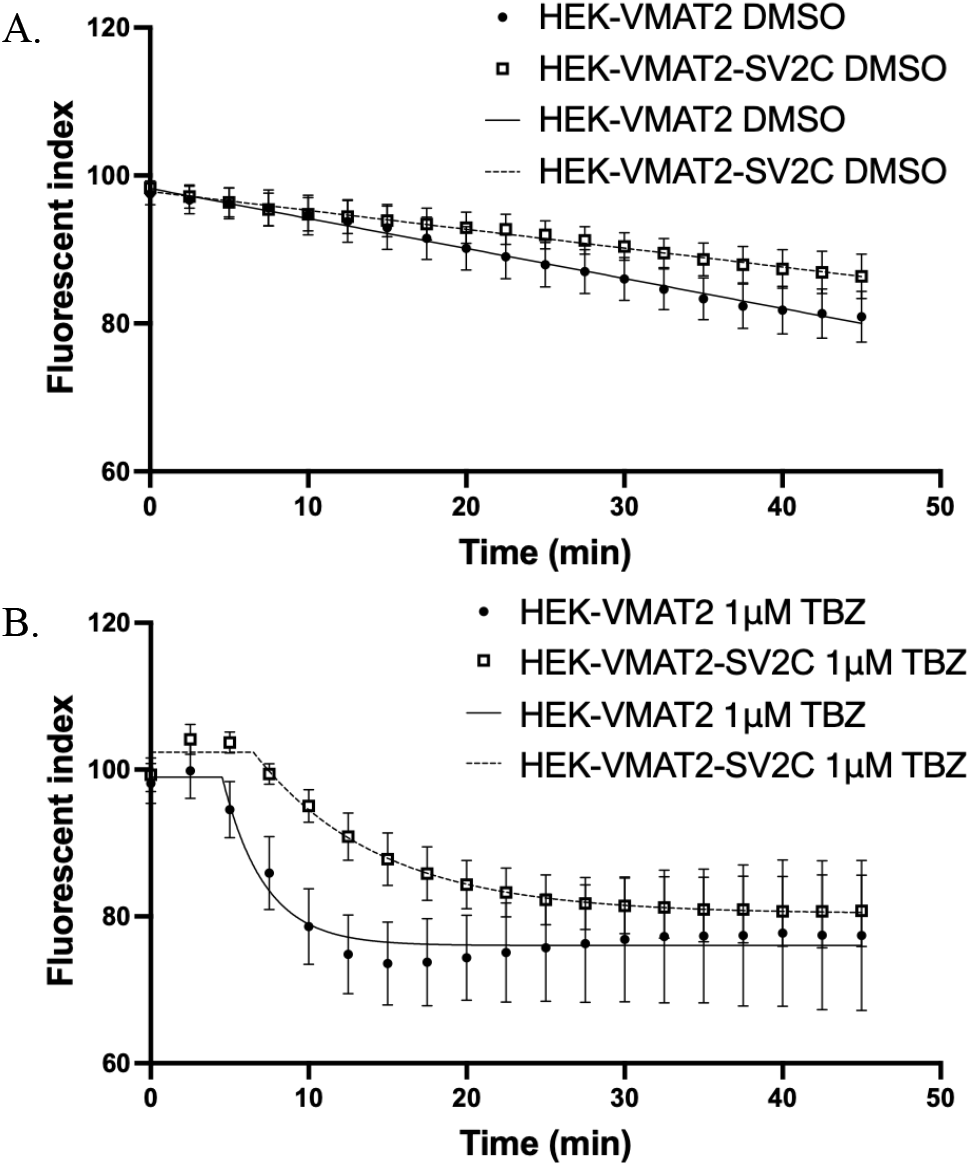
SV2C enhances retention of the dopamine analogue FFN206. FFN206 retention was measured in HEK-VMAT2 and HEK-VMAT2-SV2C cells seeded on 96-well plates. Following a scan for baseline fluorescence, control (DMSO) or tetrabenazine (1μM) was added and the plate was scanned for 45m in 2.5m intervals. Each well was normalized to its own baseline fluorescent value and values are plotted as percent of baseline. Each experiment had 12 experimental replicates per cell and treatment combination, with 10 DMSO experimental replicates and 4 tetrabenazine experimental replicates. DMSO was analyzed using linear regression and TBZ was analyzed by performing non-linear regression single-phase decay. Regressions were compared and determined that line of best fit was significantly different for each cell line in the DMSO (F(1, 376) = 6.998, p = 0.0085) and TBZ condition (F(4, 144) = 25.15, p < 0.0001).

**Figure 4.**
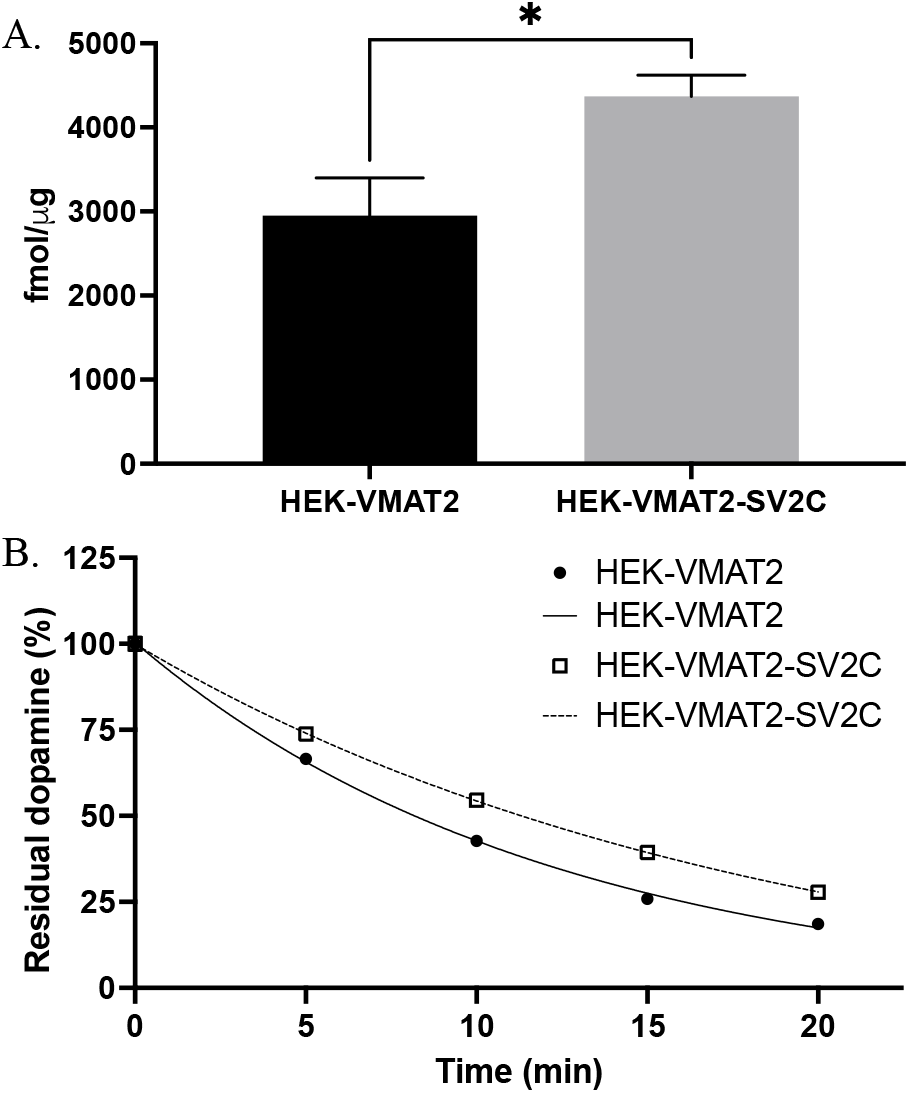
SV2C increases vesicular [^3^H]-dopamine uptake and retention in isolated vesicles. A. Vesicles were isolated from HEK-VMAT2 and HEK-VMAT2-SV2C cells before undergoing radiolabeled dopamine uptake assay. Vesicles isolated from HEK-VMAT2 cells averaged 2951.19fmol/ug of tritiated dopamine uptake compared to HEK-VMAT2-SV2C cells which averaged 4369.47fmol/ug of tritiated dopamine uptake. Unpaired t-test, n = 11-12, p =0.0102. B. SV2C slows vesicular leakage of [^3^H]-dopamine from isolated vesicles. Values were transformed into percent control of the average time_0_ value within cell line. Non-linear regression performed using one-phase decay to calculate half-life values for each cell line. HEK-VMAT2 t_1/2_ = 8.4m; n = 11-12 for each time-point. HEK-VMAT2-SV2C t_1/2_ = 12.7m; n = 11-12 for each time-point. Comparison of non-linear regressions determined the curves of best fit were significantly different for each cell line (F(3, 110) = 50.24, p <0.0001).

### Animals

Male mice (5-7 mo) were used for all 1-methyl-4-phenyl-1,2,3,6-tetrahydropyridine (MPTP) experiments due to reported sex differences and estrogen-mediated protection against MPTP intoxication.^46^ Animals were maintained on a 12/12 light/dark cycle and given food and water *ad libitum*. SV2C-KO mice were reported previously.^36^ SV2C-KO mice were generated from embryonic stem cells (ESCs) with a European Conditional Mouse Mutagenesis Program (EUCOMM) “knockout first allele” construct targeting the Sv2c gene (Sv2c^tm1a(EUCOMM)Wtsi^). The construct contained a *LacZ*/neomycin resistance cassette flanked by *FRT* sites inserted into the *Sv2c* gene. ESCs were derived from C57BL/6N-A/a mice and implanted into C57BL/6J (JAX strain #000664, RRID:IMSR_JAX:000664) female mice. Chimeric animals were crossed with a line globally expressing Flp-recombinase on a mixed C57BL/6J-129S4 background (B6.129S4-*Gt(ROSA)26Sor*^*tm2(FLP*)Sor*^/J; JAX strain # 012930, RRID:iMSR_JAX:012930) to excise the cassette, resulting in a line of mice containing a floxed exon 2 of the *Sv2c* gene. These mice were then crossed with a line containing a nestin-driven Cre-recombinase on a C57BL/6J background (B6.Cg-Tg(Nes-cre)1Kln/J, JAX strain # 003771, RRID:IMSR_JAX:003771) in order to achieve a pan-neuronal knockout of SV2C.

### MPTP injections

1-methyl-4-phenyl-1,2,3,6-tetrahydropyridine (MPTP) (Sigma or MedChemExpress) was administered according to a 5 × 20mg/kg dosing paradigm. Mice were assigned to MPTP or saline treatment randomly, and the experimenter was blinded to mouse genotype during treatment. Mice were weighed prior to MPTP administration and injected (s.c.) with 20mg/kg MPTP (freebase) or an equivalent volume of saline (control) once per day for five days, with an intrainjection interval of 24hr. The lesion was allowed to stabilize for 21 days following the final injection. Mice were sacrificed by rapid decapitation. Brains were removed, dissected, and flash frozen (for immunoblot) or post fixed in 4% (w/v) paraformaldehyde (for immunohistochemistry). Due to supply issues, data reported in Supplemental Table 1 was acquired from experiments using MPTP sourced from Sigma, and the representative immunohistochemistry images in Figure 7 and data reported in Supplemental Table 2 was acquired from experiments using MPTP sourced from MedChemExpress. A three-way ANOVA to assess main effects and interactions of MPTP source, treatment, and SV2C genotype indicated no significant main effect (cell loss: F(1,4.17), p=0.05; striatal TH: F(1,0.59), p=0.45) or interaction with other factors (cell loss: Treatment*Source: F(1,1.08), p=0.31; cell loss: Genotype*Treatment*Source: F(1,2.35), p=0.14; striatal TH: Treatment*Source: F(1,1.24), p=0.27; cell loss: Genotype*Treatment*Source: F(1,3.46), p=0.07) of MPTP source on neurotoxicity, thus stereological counts and striatal TH expression were compiled into analysis in Figure 8.

### Immunohistochemistry

Immunohistochemistry was performed as described previously described.^36^ Briefly, brains were sectioned to 40μm. Sections underwent antigen retrieval (70ºC citra buffer (Biogenix) for 1 hour) and endogenous peroxidase was quenched with 10% hydrogen peroxide. Nonspecific antibody binding was blocked with 10% normal horse serum in PBS with 0.2% Triton X-100. Tissue was incubated in primary antibody overnight at 4ºC with gentle agitation. Sections were then incubated in biotinylated secondary antibody at room temperature for 1 hour. Signal was enhanced with avidin-biotin complex (Vector) and visualized with 3-3’-diaminobenzidine (DAB). The DAB reaction was stopped with PBS. Sections were mounted to slides and stained with cresyl violet (0.1% w/v aqueous, Poly Scientific). Sections were destained with 0.1% acetic acid in 95% ethanol, followed by dehydration in ethanol and lipid-clearing in xylenes. Rabbit anti-TH (1:1,000 (Millipore AB152)).

### Stereology

Stereological cell counting was performed as described previously using StereoInvestigator software.^45,47,48^ Experimenters were blinded to mouse genotype and treatment throughout the experiment, including during immunohistochemistry and stereological analysis. Samples were randomized during staining and stereological analysis. Every fourth SNpc section was counted using the optical fractionator method. Counting frames of 50μm X 50μm on a 120μm X 120μm counting grid were sampled. Cell counts were weighted based on manually measured section thickness. Only counts with a Gunderson coefficient of error (m=0) of <0.10 were included for analysis.

### Statistics

Graphical representations and statistical analysis was performed in GraphPad Prism 9.

## Results

### Creation of double-stable HEK-VMAT2-SV2C cell line

To investigate the effect that SV2C has on vesicular dynamics, we created a double-stable cell line (HEK-VMAT2-SV2C) by introducing human SV2C into HEK293 cells stably expressing human vesicular monoamine transporter 2 (VMAT2; HEK-VMAT2) previously utilized in Black *et al*. 2021.^44^ The expression of SV2C in HEK-VMAT2-SV2C cells is detectable by Western blot with HEK-VMAT2 cells demonstrating no endogenous expression of SV2C, and HEK-VMAT2-SV2C cells demonstrating robust SV2C expression (Figure 1A). When visualized with immunocytochemistry, SV2C co-localizes with VMAT2 with punctate staining on subcellular compartments (Figure 1B,C).

### Effect of SV2C on FFN206 uptake in HEK-VMAT2 cells

Our previously developed *in vitro* assay allows for visualization of vesicular dynamics utilizing the fluorescent dopamine analogue and VMAT2 substrate fluorescent false neurotransmitter 206 (FFN206).^44^ In this assay, HEK-VMAT2 cells are incubated with FFN206, which accumulates within vesicular compartments via VMAT2-mediated transport. The accumulation of FFN206 can be blocked with VMAT2 inhibitors and by dissipating the proton gradient VMAT2 requires to load substrate within subcellular compartments.^44^ When FFN206 accumulates within vesicular compartments, the fluorescent signal is concentrated allowing for visualization and measurement of the substrate by microscopy and plate reader analysis. To investigate the effect that SV2C has on the vesicular sequestration of FFN206, we measured the uptake of FFN206 in HEK-VMAT2-SV2C cells compared to HEK-VMAT2 cells.

The addition of SV2C to HEK-VMAT2 cells results in enhanced uptake of FFN206 as visualized by microscopy (Figure 2A). Quantification of uptake performed in 96-well plates on a plate reader detected a 33.05% increase in FFN206 fluorescence in HEK-VMAT2-SV2C cells compared to HEK-VMAT2 cells (Two-way ANOVA with Šídák’s multiple comparisons test, n = 7 experimental replicates, p <0.005, Figure 2C). The uptake of FFN206 is inhibited in both HEK-VMAT2 and HEK-VMAT2-SV2C cells by pre-treating the cells with a saturating dose (1μM) of the VMAT2 inhibitor tetrabenazine (TBZ) before FFN206 application. (Two-way ANOVA with Šídák’s multiple comparisons test, n = 7 experimental replicates, p = 0.83, Figure 2B). We have previously reported the IC_50_ of TBZ in HEK-VMAT2 cells to be 73 nM.^44^ A TBZ dose-response to determine whether HEK-VMAT2-SV2C cells are resistant to TBZ-induced inhibition revealed a significant difference in IC_50_ values, with HEK-VMAT2 cells demonstrating an IC_50_ of 135 nM, and HEK-VMAT2-SV2C cells demonstrating an IC_50_ value of 275 nM (non-linear regression [inhibitor] vs. normalized response, n = 4-8 experimental replicates for each dose, p < 0.05, Figure 2C).

### Effect of SV2C on FFN206 retention in HEK-VMAT2 cells

To investigate the effect of SV2C on the retention of VMAT2 substrates in vesicles, we developed a novel retention assay to measure FFN206 fluorescence over time. Cells were incubated with FFN206 for 1 hour to allow for VMAT2-mediated uptake within vesicular compartments. Following incubation, the FFN206 solution was removed from the cells and a baseline measurement of fluorescence was obtained. The fluorescence was continually measured in 2.5m intervals for 45 minutes and expressed as a percent of baseline fluorescence as a representation of the retention of FFN206 over time.

We found that while FFN206 fluorescence linearly decays over time for both HEK-VMAT2 and HEK-VMAT2-SV2C cells (Figure 3A), HEK-VMAT2-SV2C cells demonstrate a modest but significant protection against loss of fluorescence over time based on linear regression. The slope of HEK-VMAT2 cells in a DMSO condition equals -0.41, representing a 0.41% loss of fluorescence per minute, whereas the slope for HEK-VMAT2-SV2C cells in a DMSO condition is -0.25, representing a 0.25% loss of fluorescence per minute (linear regression, n = 10 experimental replicates, p <0.01, Figure 3A). Inhibiting VMAT2 with TBZ prevents the reuptake of FFN206 by VMAT2 from the cytosol into the subcellular compartment, thus ensuring that the fluorescent values being measured were representative of the pool of FFN206 being retained within vesicles. To compare the rate of decay of fluorescence in the TBZ condition, non-linear regression with single-phase decay was performed. HEK-VMAT2 cells demonstrate a rapid loss in fluorescence following TBZ treatment, whereas the HEK-VMAT2-SV2C cells show moderate protection against this loss of fluorescence. Analysis of non-linear regressions determined the decay curves are significantly different for HEK-VMAT2 cells treated with TBZ compared to HEK-VMAT2-SV2C cells treated with TBZ (non-linear regression with one-phase exponential decay, n = 4 experimental replicates, p < 0.0001, Figure 3B).

### Effect of SV2C on [^3^H]-dopamine uptake and retention in vesicles derived from HEK-VMAT2 cells

Although the fluorescent false neurotransmitters, including FFN206, were designed to be fluorescent substrates for VMAT2,^49^ we investigated whether these findings would be replicated with dopamine as the substrate utilizing tritiated dopamine ([^3^H]-dopamine) in radiolabeled uptake and retention assays in vesicles isolated from HEK-VMAT2 and HEK-VMAT2-SV2C cells. The addition of SV2C to HEK-VMAT2 cells resulted in a 46.7% increase in uptake of [^3^H]-dopamine compared to HEK-VMAT2 cells lacking SV2C (Unpaired t-test, n = 11-12, p < 0.05, Figure 4A). To measure how much [^3^H]-dopamine was retained within vesicles over time, vesicles were incubated with [^3^H]-dopamine before the VMAT2 inhibitor reserpine (10μM) was applied to the vesicles. This effectively froze the transporter, allowing us to compare a time course of vesicular leak to uninhibited vesicles. The remaining [^3^H]-dopamine was measured at five time-points post-reserpine application (0m, 5m, 10m, 15m, and 20m) to determine the rate of [^3^H]-dopamine leakage over time. By comparing the half-life calculated from the rate of decay in each cell line, it was determined that HEK-VMAT2-SV2C cells had a slower rate of leakage with a t_1/2_ = 12.7m compared to a t_1/2_ = 8.4m for HEK-VMAT2 cells (non-linear regression with one-phase exponential decay, n = 11-12 for each time-point, p <0.0001, Figure 4B).

### Effect of SV2C on [^3^H]-dopamine uptake and retention in brain-derived vesicles from WT and SV2C-KO mice

Because the subcellular compartments on which VMAT2 and SV2C localize in HEK293 cells are not synaptic vesicles per se, we next determined whether the results of the radiolabeled uptake and retention assays were replicated in synaptic vesicles derived from mouse brain tissue. Vesicles were isolated from brain homogenate of wild-type and SV2C-KO animals and radiolabeled dopamine uptake and retention assays were performed. Although SV2C-KO animals showed no difference in [^3^H]-dopamine uptake compared to wild-type littermate controls as determined by a dose-response measure of vesicular capacity (Supplemental Figure 1), vesicles derived from SV2C-KO animals had significantly reduced dopamine retention with a t_1/2_ = 1.98m compared to vesicles derived from wild-type littermate controls, which had a t_1/2_ = 3.32m (non-linear regression with one-phase exponential decay, n = 3-4 for each time-point, p <0.005, Figure 5).

**Figure 5.**
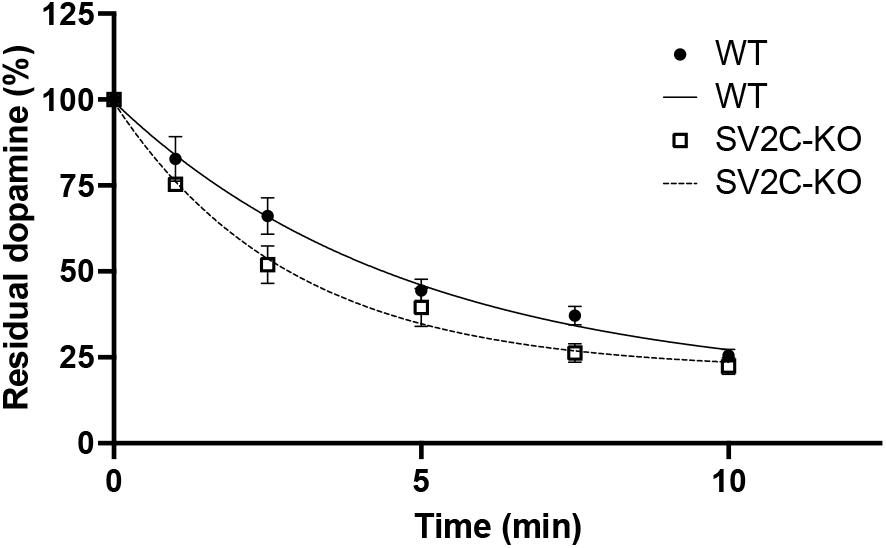
Genetic ablation of SV2C increases vesicular leakage of [^3^H]-dopamine in brain-derived vesicles. Values were transformed into percent control of the average time_0_ value within mouse line. Non-linear regression performed using one-phase decay to calculate half-life values for each mouse line. WT t_1/2_ = 3.3m; n = 3-4 for each time-point. SV2C-KOt_1/2_ = 2.0m; n = 3-4 for each time-point. Comparison of non-linear regressions determined the curves of best fit were significantly different for each mouse line (F(3, 39) = 5.546, p =0.0029).

### Effect of SV2C on [^3^H]-MPP^+^ uptake and retention in brain-derived vesicles from WT and SV2C-KO mice

The expression of VMAT2 is known to mediate dopamine neuron health by sequestering both dopamine, and the exogenous neurotoxicant and VMAT2 substrate MPP^+^.^11,12,47^ Following our results indicating that SV2C mediates the vesicular storage of dopamine and the dopamine analogue FFN206, we next set out to determine whether SV2C may play a similar role in mediating the vesicular storage of MPP^+^. Radiolabeled uptake and retention assays were performed in vesicles isolated from HEK-VMAT2 and HEK-VMAT2-SV2C cells using radiolabeled MPP^+^ ([^3^H]-MPP^+^). There was a 44.6% increase in the uptake of [^3^H]-MPP^+^ in HEK-VMAT2-SV2C cells compared to HEK-VMAT2 cells (Unpaired t-test, n = 12, p < 0.05, Figure 6A). In the retention assay, the half-life calculated from the rate of decay in each cell line demonstrated HEK-VMAT2-SV2C cells had a slower rate of leakage with a t_1/2_ = 4.3m compared to a t_1/2_ = 3.5m for HEK-VMAT2 cells (non-linear regression with one-phase exponential decay, n = 10-12 for each time-point, p <0.0005) (Figure 6B). These data indicate that in addition to regulating the vesicular storage of dopamine, SV2C may confer neuroprotection by mediating vesicular storage of exogenous neurotoxicants.

**Figure 6.**
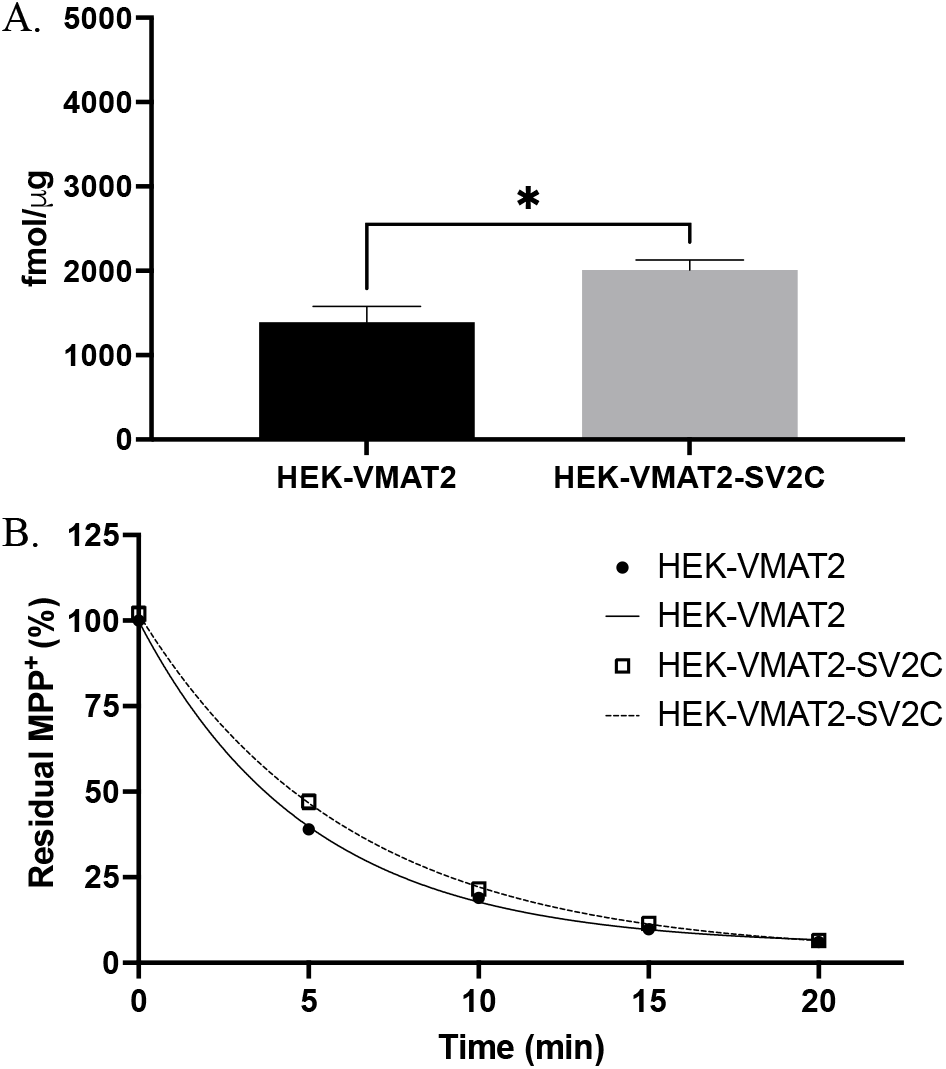
SV2C increases vesicular uptake and retention of [^3^H]-MPP^+^. A. Vesicles were isolated from HEK-VMAT2 and HEK-VMAT2-SV2C cells before undergoing uptake assay. Vesicles isolated from HEK-VMAT2 cells averaged 1390.31fmol/ug of tritiated MPP^+^ uptake compared to HEK-VMAT2-SV2C cells which averaged 2010.14fmol/ug of tritiated MPP^+^ uptake. Unpaired t-test, n = 12, p =0.0106. B. SV2C increases vesicular retention of [^3^H]-MPP^+^. Values were transformed into percent control of the average time_0_ value within cell line. Non-linear regression performed using one-phase decay to calculate half-life values for each cell line. HEK-VMAT2 t_1/2_ = 3.5m; n = 10-12 for each time-point. HEK-VMAT2-SV2C t_1/2_ = 4.3m; n = 10-12 for each time-point. Comparison of non-linear regressions determined the curves of best fit were significantly different for each cell line (F(3, 107) = 6.712, p =0.0003).

**Figure 7.**
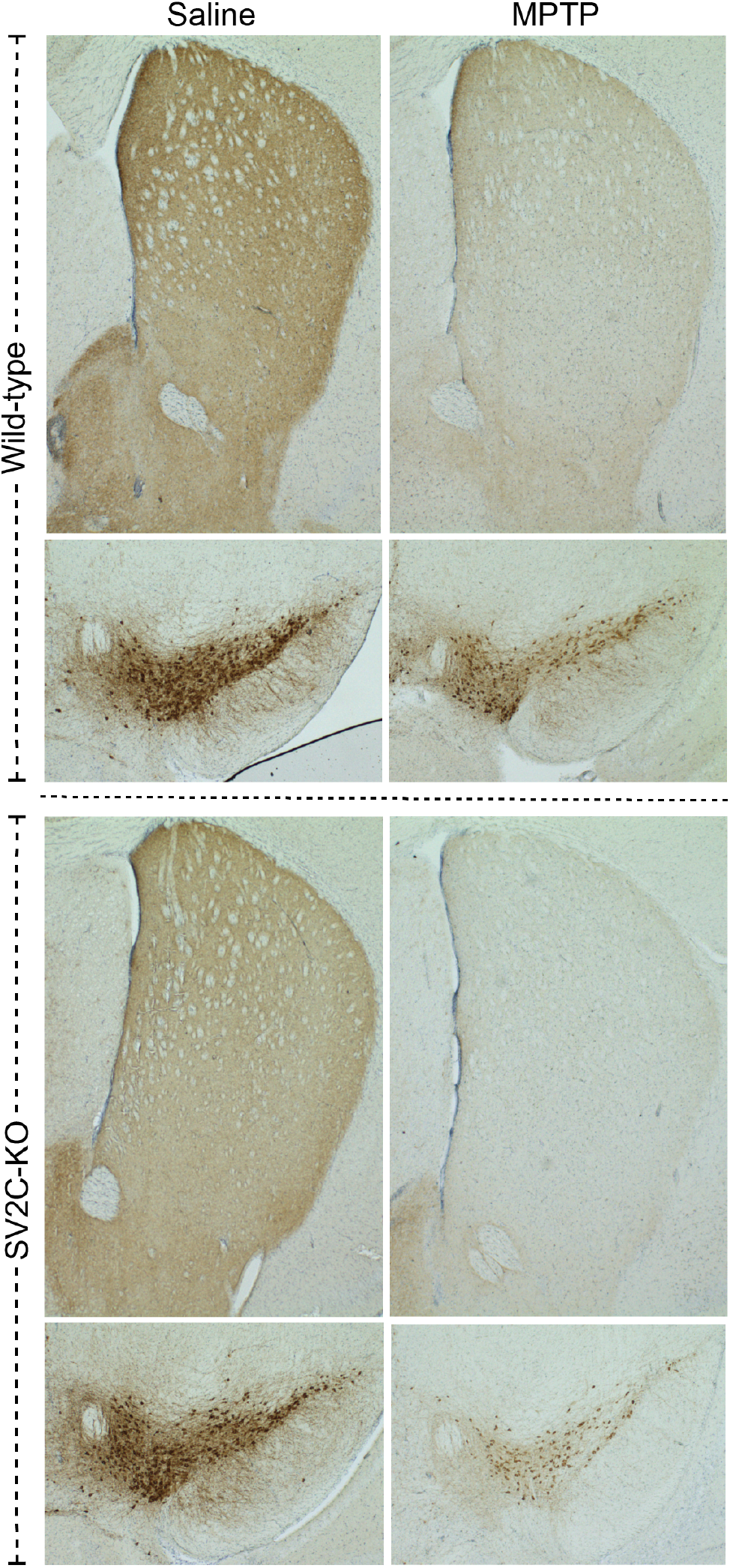
Genetic ablation of SV2C enhances dopamine vulnerability to MPTP. Representative TH immunohistochemistry of the striatum and midbrain of wild-type and SV2C-KO animals treated with saline or MPTP demonstrates that the lesion to the basal ganglia is enhanced in both the dorsal striatum and substantia nigra of SV2C-KO animals (bottom) as compared to wild-type controls (top).

**Figure 8.**
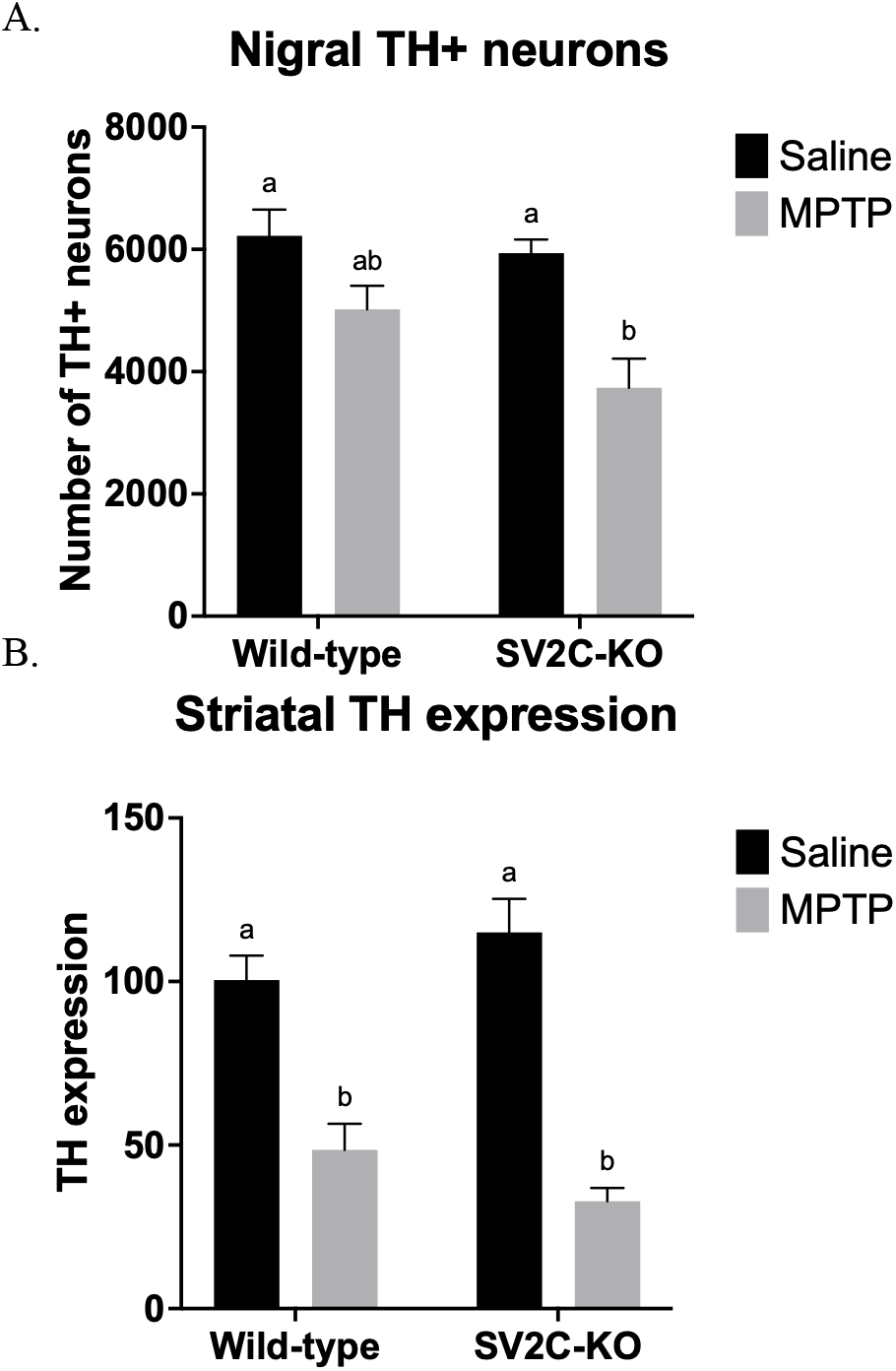
Genetic ablation of SV2C results in loss of nigral TH+ neurons following mild MPTP treatment regimen. A. Quantification of intact dopaminergic cells of the SNc using unbiased stereological cell counting confirms a significant loss dopaminergic nigral cells in SV2C-KO, but not WT animals following MPTP. Nigral TH+ neuronal counts: Two-way ANOVA with Tukey’s multiple comparisons test. Effect of genotype (F(1, 33) = 3.487, p = 0.0707); effect of treatment F(1, 33) = 16.47, p = 0.0003); effect of interaction (F(1, 33) = 1.429, p = 0.2405). n = 6-12 animals per genotype:treatment group. WT:saline vs. WT:MPTP p =0.2713; WT:saline vs. SV2C-KO:MPTP p =0.0017; SV2C-KO:saline vs. SV2C-KO:MPTP p =0.0011; WT:MPTP vs. SV2C-KO:MPTP p =0.1055. B. Quantitation of TH expression from striatal lysate by Western blot shows significant loss of TH expression following MPTP compared to within genotype control. Striatal TH expression: Two-way ANOVA with Tukey’s multiple comparisons test. Effect of genotype (F(1, 37) = 0.00526, p = 0.9246); effect of treatment F(1, 37) = 74.39, p < 0.0001); effect of interaction (F(1, 37) = 3.795, p = 0.0590). n = 6-12 animals per genotype:treatment group. WT:saline vs. WT:MPTP p =0.0007; WT:saline vs. SV2C-KO:MPTP p <0.0001; WT:MPTP vs. SV2C-KO:saline p <0.0001; SV2C-KO:saline vs. SV2C-KO:MPTP p <0.0001.

### Protective effect of SV2C on MPTP-induced neurotoxicity in WT and SV2C-KO mice

To evaluate whether ablation of SV2C results in heightened vulnerability to MPTP and its active metabolite MPP^+^, we administered a 5×20mg/kg dose of MPTP (Sigma or MedChemExpress) consisting of five injections (s.c.) over five days with an inter-injection interval of 24hr in SV2C knock-out (SV2C-KO) mice and wild-type littermate controls. Immunohistochemistry of tyrosine hydroxylase (TH) revealed enhanced loss of immunoreactivity in the striatum and substantia nigra (Figure 7). To determine if there was an exacerbation of dopaminergic cell body loss following MPTP, we performed stereological cell counting of TH-positive cells in the substantia nigra pars compacta (SNc) (Figure 8a). Baseline dopaminergic cell count was equivalent between SV2C-KO and WT animals (WT: 6221 cells ± 1049; SV2C-KO: 5939 cells ± 698, p =0.97). Although there was a trend toward fewer neurons in WT animals following MPTP administration, this did not reach significance (p =0.27); however, SV2C-KO animals demonstrated a significant loss of 37.1% fewer TH-positive neurons in the SNc following MPTP (p <0.005) (Two-way ANOVA with Tukey’s multiple comparisons test, n = 6-12.) Quantification of TH expression in the striatum performed by Western blot revealed a significant loss of TH expression in both wild-type and SV2C-KO controls exposed to MPTP in comparison to within genotype controls (Figure 8b, Two-way ANOVA with Tukey’s multiple comparisons test, n = 6-12. WT:saline vs. WT:MPTP p <0.001; SV2C-KO:saline vs. SV2C-KO:MPTP p <0.0001).

## Discussion

Our previous *in vivo* work in SV2C knockout mice implicated SV2C as a modifier of vesicular dopamine function.^36^ We have previously shown that HEK293 cells expressing human VMAT2 accumulate FFN206 within subcellular components that colocalize with VMAT2.^44^ Here, we introduce HEK293 cells that stably express both human VMAT2 and human SV2C (Figure 1), which show colocalization of VMAT2 and SV2C on subcellular compartments and enhanced accumulation of FFN206 (Figure 2) and radiolabeled dopamine (Figure 4). Additionally, we present the first evidence that SV2C exerts its effects on the vesicular storage of dopamine and the dopamine analogues FFN206 and MPP^+^ by enhancing total vesicular uptake and promoting vesicular retention.

The use of the fluorescent VMAT2 substrate FFN206 further allows for real-time visualization and monitoring of vesicular dynamics, which we utilized to develop a novel assay to monitor vesicular retention. Because vesicles are inherently permeant or leaky, it is possible to measure the retention of substrates within vesicles over time. FFN206 fluorescence is visualizable and measurable only when it is accumulated within subcellular compartments; thus, diffuse and dilute FFN206 is not detected by fluorescent plate reader or microscopy. This property of FFN206 allows for calculation of vesicular retention by measuring the fluorescence of accumulated FFN206 at baseline and tracking fluorescent values over time. We developed a plate reader-based assay using cells seeded on 96-well plates that measures the fluorescence of each well over time. Values can be corrected for background fluorescence by including cells receiving the same treatments (e.g., DMSO and tetrabenazine) that were not incubated with FFN206 each time the assay is run and performing a background subtraction using the average values calculated from these wells. This background subtraction is applied at each time-point that the plate is scanned over the course of the assay and the resulting fluorescent value can be expressed as a percent of its baseline (e.g., time-point 0m) fluorescence. Plotting the percent of baseline fluorescence over time allows for regression analysis to measure the loss of fluorescence over time.

Measuring vesicular retention of FFN206 identified that HEK293 cells expressing both human VMAT2 and human SV2C (HEK-VMAT2-SV2C) retain FFN206 within vesicles better than cells only expressing VMAT2 (HEK-VMAT2) (Figure 3). Although this is a modest effect in cells treated with DMSO as a control treatment, it is made more apparent by adding the VMAT2 inhibitor TBZ at a saturating concentration (1μM). The vesicular packaging of substrates is a dynamic process involving leakage and reuptake of vesicular contents, and active transport of components in and out of the vesicles to maintain vesicular pH and achieve a functional equilibrium. While measuring FFN206 fluorescence over time, in the DMSO condition, it is possible that as FFN206 leaks out of the vesicle it diffuses beyond the local vesicular environment and is unable to be resequestered by VMAT2 or otherwise undergoes degradation. By adding TBZ, the resequestration by VMAT2 of FFN206 that leaks out of the vesicle can be prevented. As an added consequence, the application of TBZ also precludes the active export of FFN206 out of the vesicle by VMAT2. Thus, the observed decay in fluorescence over time is due to FFN206 leaking out of the vesicle in a non-VMAT2 dependent manner and is indicative of how much FFN206 remains sequestered within vesicles.

Although FFN206 was designed as a dopamine analogue and substrate for VMAT2, we sought to confirm that the effect of SV2C on vesicular dynamics using FFN206 were replicated using dopamine directly. Vesicles isolated from HEK293 cells expressing human VMAT2 or both human VMAT2 and SV2C underwent uptake assays using radiolabeled dopamine ([^3^H]-dopamine) and demonstrated enhanced uptake and retention of [^3^H]-dopamine (Figure 4). Retention, or leakage, assays were performed in a similar manner to FFN206 retention assays where vesicles were incubated with [^3^H]-dopamine before sequestration was blocked with the VMAT2 inhibitors TBZ or reserpine and the amount of [^3^H]-dopamine remaining within the vesicles was measured at multiple time-points. However, because HEK293 cells are non-neuronal and do not contain all of the same factors as neurons, we next determined if these functions occur in vesicles isolated from mouse brain.

While uptake of FFN206 is increased in whole HEK-VMAT2-SV2C cells, and uptake of and [^3^H]-dopamine is increased in vesicles derived from HEK-VMAT2-SV2C cells, vesicles isolated from mice with genetic ablation of SV2C (SV2C-KO) do not show a difference in total vesicular uptake of [^3^H]-dopamine (Supplemental Figure 1). This may be due to differences in the nature of the compartments within HEK293 cells on which VMAT2 and SV2C localize compared to the composition of synaptic vesicles. For example, in HEK293 cells, these compartments may have increased capacity for the total amount of substrate uptake based on size or number of copies of VMAT2 expressed on each compartment, or increased uptake due to increased pool of protons within the compartments providing the proton-motive force by which VMAT2 loads substrates. Despite there being no difference in the baseline amount of [^3^H]-dopamine uptake, synaptic vesicles derived from the brain of SV2C-KO mice displayed enhanced rate of [^3^H]-dopamine leak compared to those from wild-type animals (Figure 5).

HEK-VMAT2-SV2C cells appear to be resistant to the effects of the VMAT2 inhibitor tetrabenazine (TBZ) when measuring FFN206 uptake (Figure 2C). It is possible that the presence of SV2C renders VMAT2 more functional in taking up dopamine or more resistant to the inhibitory effects of TBZ. Furthermore, it is possible that SV2C enhances vesicular dopamine uptake and retention through direct transport of dopamine itself. However, if SV2C could transport dopamine, we would expect to see a significant difference in FFN206 uptake in cells expressing both VMAT2 and SV2C compared to cells only expressing VMAT2 when uptake by VMAT2 is fully inhibited (e.g., with pre-treatment of >1μM TBZ) (Figure 2C). Further experiments will be necessary to determine whether the shift in inhibition is due to an SV2C-mediated decrease of TBZ binding or efficacy at VMAT2. However, the results may also be a result of the initial non-specific accumulation within HEK293 cells observed with FFN206, which can enter HEK293 cells that do not express the plasmalemmal dopamine transporter (DAT). Initial accumulation within the cell is not dependent on dopamine transporters (e.g., DAT and VMAT2); however, once FFN206 has reached diffuse equilibrium throughout the cell, its accumulation within vesicles is dependent upon VMAT2. Thus, SV2C may be acting to retain the FFN206 that has accumulated within vesicles in a non-VMAT2 dependent manner, thereby resisting the effects of TBZ.

It has been well-established that MPTP toxicity can regulated by dopaminergic synaptic vesicles.^11,25,50-54^ In animal models MPTP administered systemically results in dopaminergic neuron degeneration in the brain due to the toxic metabolite MPP^+^ acting as a substrate for the plasmalemmal dopamine transporter (DAT), allowing for accumulation within dopaminergic neurons. When dopamine vesicles have a greater capacity to store dopamine and sequester toxicants from the cytosol, neurons are resistant to degeneration, and when this process is impaired, neurons are vulnerable to enhanced cellular damage and death. This has been demonstrated repeatedly by our laboratory in models of varying VMAT2 expression.^47,51^ In fact, VMAT2 was originally identified for its role in sequestering the active metabolite of MPTP, MPP^+^, and protecting against MPTP toxicity.^55^

Our evidence from experiments evaluating the effect of SV2C on FFN206 and [^3^H]-dopamine uptake and retention suggests that SV2C may similarly interact with the neurotoxicant, and VMAT2 substrate, MPP^+^. Here, we provide the first evidence that another vesicular protein, SV2C, may promote the vesicular retention of the neurotoxicant MPP^+^ and mediate the toxic effects of MPTP. By isolating vesicles from HEK293 cells expressing VMAT2 and SV2C, we were able to directly interrogate the effect SV2C has on uptake and retention of radiolabeled MPP^+^ ([^3^H]-MPP^+^). Similar to the results from [^3^H]-dopamine experiments, we found that cells expressing both VMAT2 and SV2C had enhanced uptake and retention of [^3^H]-MPP^+^ (Figure 6). Interestingly, while the extent of increased uptake in vesicles from cells containing both VMAT2 and SV2C was comparable, with a 48.1% increase in [^3^H]-dopamine uptake and a 44.6% increase in [^3^H]-MPP^+^ uptake, the raw values of uptake were much higher for [^3^H]-dopamine than [^3^H]-MPP^+^, with uptake values of 4369 fmol/μg protein and 2010.14 fmol/μg protein respectively. Furthermore, the rate of leakage is different between [^3^H]-dopamine and [^3^H]-MPP^+^, with a t_1/2_ equal to 12.7m and 4.3m respectively. Other studies have reported similar affinities of VMAT2 for dopamine and MPP^+^.^56^ The design of the experiments conducted in this study provide interesting data warranting deeper investigation in future studies looking at competition between dopamine and MPP^+^ as substrates.

Following the data from radiolabeled uptake and retention experiments, we next investigated the biological implications *in vivo* with experiments exposing wild-type and SV2C-KO animals to MPTP. As a result, we present data that showing that SV2C is protective against chemically-induced neurodegeneration. Mice with genetic ablation of SV2C demonstrate a greater effect of MPTP on nigral and striatal tyrosine hydroxylase (TH) immunoreactivity, indicating increased vulnerability (Figure 7). While there is no significant difference in striatal TH expression as determined by Western blot quantitation (Figure 8b), SV2C-KO mice treated with MPTP have significantly fewer TH-positive neuronal cell bodies in the substantia nigra compared to control treated SV2C-KO mice, whereas wild-type mice treated with MPTP do not show a significant difference in the number of TH-positive neuronal cell bodies in the substantia nigra compared to control treated wild-type mice (Figure 8a). Impairment of vesicular function as a result of SV2C-KO may result in higher cytosolic concentrations of dopamine and/or MPP^+^. Cytosolic dopamine can act as an endogenous neurotoxicant, and cytosolic MPP^+^ is free to act as a mitochondrial complex I inhibitor, both of which may contribute to neuron vulnerability.

Although HEK293 cells are non-neuronal and do not contain all the proteins involved in vesicular sequestration that can be found in brain tissue, experiments in these cells that display proton gradient dependent storage of dopamine in a vesicle-like compartment allow for isolation of the factors involved to include only the effect of VMAT2 and SV2C. The comparable results seen with multiple VMAT2 substrates (e.g., FFN206, [^3^H]-dopamine, and [^3^H]-MPP^+^) in multiple experimental systems (e.g., whole HEK293 cells, isolated vesicles from HEK293 cells, and brain derived vesicles from mouse brain) provides additional evidence, additionally supported by the work of other groups^49^, in support of using FFN206 as an appropriate proxy for evaluating dopamine dynamics.^57^ This validation can be applied to future experiments to reduce animal use, experiment cost, and exposure to radiolabeled compounds by instead initially performing experiments in cells with FFN206. There are several unanswered questions with regard to the function of SV2C. For instance, SV2 proteins are highly glycosylated and the intraluminal loops of SV2s are thought to comprise the intra-vesicular proteoglycan “gel” matrix, which can be visualized by electron microscopy.^27^ The proteoglycan matrix within vesicles has been demonstrated as capable of directly adsorbing ATP as visualized by atomic force microscopy and is hypothesized to regulate the release of transmitter molecules into the synaptic cleft upon endocytosis.^27,58-60^ In future experiments, the role of glycosylation in SV2C function can be evaluated with site directed mutations of the glycosylation sites on SV2C.

Overall, these data further support the emergence of SV2C as a relevant player in dopamine neuron and vesicle function. We establish SV2C as a mediator of MPTP toxicity, and suggest that SV2C function and expression is inversely correlated with vulnerability to degeneration. These data point to a role for SV2C in mediating chemically-induced dopaminergic degeneration, and future studies will investigate whether this neuroprotective function of SV2C may extend to additional models of cell death. Additionally, these data may suggest that enhanced SV2C could be neuroprotective, which could have significant implications for the development of Parkinson’s disease therapeutics.

## Acknowledgments

We would like to acknowledge David Sulzer for his help with optimizing fluorescent false neurotransmitter assays and Hae Jung Chung for her technical assistance with western blot protocols.

## Ethical statement

All procedures were conducted in accordance with the National Institutes of Health Guide for Care and Use of Laboratory Animals (Committee on Care and Use of Laboratory Animals (1996) Guide for the Care and Use of Laboratory Animals (Natl Inst Health, Bethesda), DHHS Publ No (NIH) 85-23.) and were previously approved by the Institutional Animal Care and Use Committee at Emory University.

## Data availability statement

The data that support the findings of this study are openly available in dryad at https://datadryad.org/stash/share/57AMYLbCm-p7yo0mD4weTg0bi5Ef9tFf5_QdjEjBTRY.

