## Supplemental Figures for "Synaptic vesicle glycoprotein 2C enhances vesicular storage of dopamine and counters dopaminergic toxicity"

Supplemental Figure 1. Genetic ablation of SV2C does not affect vesicular capacity of [$^3$H]-dopamine uptake in brain-derived vesicles

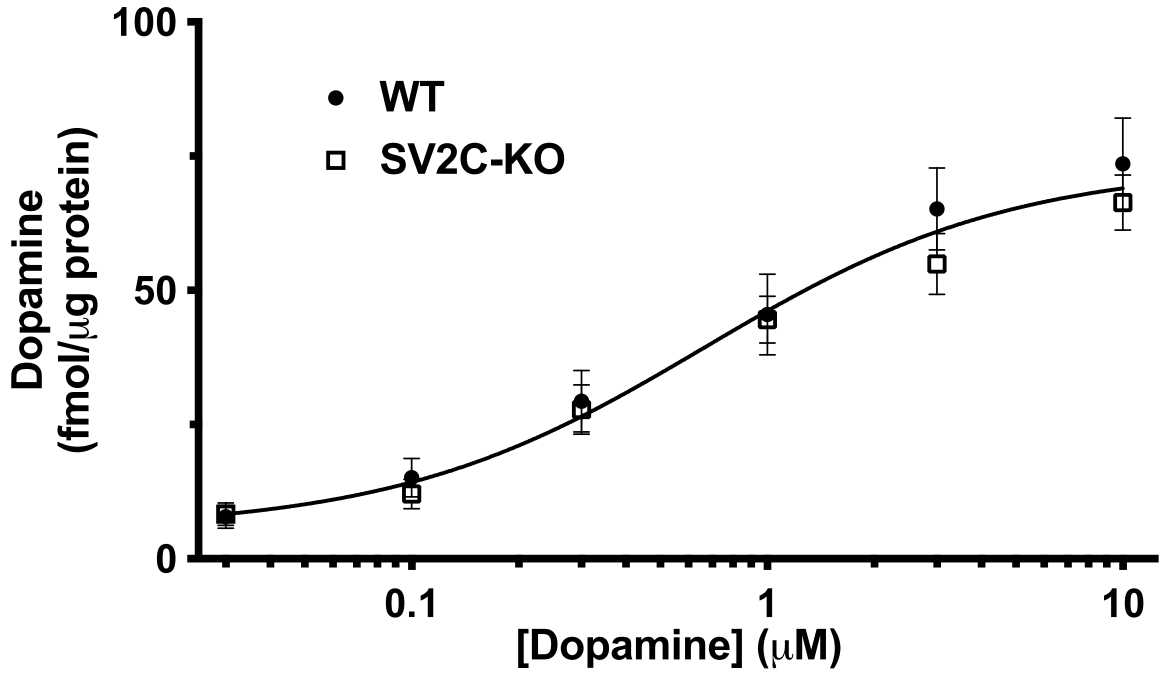


Supplemental Figure 1. Genetic ablation of SV2C does not affect vesicular capacity of [$^3$H]-dopamine uptake in brain-derived vesicles. Radiolabeled dopamine uptake in isolate vesicles from WT and SV2C-KO animals (n = 7 animals per genotype) across multiple concentrations of dopamine (0.03μM - 10μM).

| | Wild-type | | SV2C knock-out | |
|---|---|---|---|---|
| | **Saline** | **Sigma MPTP** | **Saline** | **MPTP** |
| **Nigral TH+ neurons** | 6053 ± 364[ab] | 4182 ± 244[bc] | 6036 ± 268[a] | 4033 ± 520[bc] |
| **Striatal TH expression** | 100 ± 8.9[a] | 33.2 ± 4.9[b] | 108 ± 7.7[a] | 30 ± 4.8[b] |

Supplemental table 1. Genetic ablation of SV2C results in increased loss of nigral TH+ neurons following MPTP (Sigma) as compared to wild-type animals. Quantification of intact dopaminergic cells of the SNc using unbiased stereological cell counting confirms a significant loss dopaminergic nigral cells in SV2C-KO, but not WT animals following MPTP.

Nigral TH+ neuronal counts: Two-way ANOVA with Tukey's multiple comparisons test, effect of genotype ($F(1, 23) = 0.03027$, $p = 0.8634$); effect of treatment ($F,(1, 23) = 16.45$, $p = 0.0005$; effect of interaction ($F(1, 23) = 0.01901$, $p = 0.8915$). n = 4-10 animals per genotype:treatment group. WT:saline vs. SV2C-KO:MPTP $p = 0.0347$; WT:MPTP vs. SV2C-KO:saline $p = 0.0469$; SV2C-KO:saline vs. SV2C-KO:MPTP $p = 0.0071$.

Striatal TH expression: Two-way ANOVA with Tukey's multiple comparisons test, effect of genotype ($F(1, 23) = 0.1351$, $p = 0.7166$); effect of treatment ($F,(1, 23) = 106.3$, $p =< 0.0001$; effect of interaction ($F(1, 23) = 0.6757$, $p = 0.4195$). n = 4-10 animals per genotype:treatment group. WT:saline vs. WT:MPTP $p < 0.0001$; WT:saline vs. SV2C-KO:MPTP $p < 0.0001$; WT:MPTP vs. SV2C-KO:saline $p < 0.0001$; SV2C-KO:saline vs. SV2C-KO:MPTP $p < 0.0001$.

| | Wild-type | | SV2C knock-out | |
|---|---|---|---|---|
| | **Saline** | **MedChemExpress MPTP** | **Saline** | **MPTP** |
| **Nigral TH+ neurons** | 6557 ± 1337[a] | 6070 ± 359[a] | 5554 ± 15[a] | 2260 ± 316[b] |
| **Striatal TH expression** | 101 ± 15.4[a] | 67.8 ± 11.0[a] | 100 ± 40.8[a] | 40 ± 7.1[b] |

Supplemental Table 2. Genetic ablation of SV2C results in increased loss of nigral TH+ neurons following MPTP (MedChemExpress) as compared to wild-type animals. Quantification of intact dopaminergic cells of the SNc using unbiased stereological cell counting confirms a significant loss dopaminergic nigral cells and striatal TH expression in SV2C-KO, but not WT animals following MPTP.

Nigral TH+ neuronal counts: Two-way ANOVA with Tukey's multiple comparisons test, effect of genotype ($F(1, 6) = 14.92$, $p = 0.0652$); effect of treatment (F,(1, 6) = 9.214, $p = 0.0229$; effect of interaction ($F(1, 6) = 5.075$, $p = 0.0652$). n = 2-4 animals per genotype:treatment group. WT:saline vs. SV2C-KO:MPTP $p = 0.0151$; WT:MPTP vs. SV2C-KO:MPTP $p = 0.0135$; SV2C-KO:saline vs. SV2C-KO:MPTP $p = 0.480$.

Striatal TH expression: Two-way ANOVA with Tukey's multiple comparisons test, effect of genotype ($F(1, 10) = 0.01255$, $p = 0.9130$); effect of treatment (F,(1, 10) = 12.98, $p = 0.0048$; effect of interaction ($F(1, 10) = 2.889$, $p = 0.1200$). n = 2-4 animals per genotype:treatment group. SV2C-KO:saline vs. SV2C-KO:MPTP $p = 0.0167$.
